# Global trade-offs in tree functional traits

**DOI:** 10.1101/2021.09.16.458157

**Authors:** Daniel S. Maynard, Lalasia Bialic-Murphy, Constantin M. Zohner, Colin Averill, Johan van den Hoogen, Haozhi Ma, Lidong Mo, Gabriel Reuben Smith, Isabelle Aubin, Erika Berenguer, Coline C.F. Boonman, Jane Catford, Bruno E. L. Cerabolini, Arildo S. Dias, Andrés González-Melo, Peter Hietz, Christopher H. Lusk, Akira S. Mori, Ülo Niinemets, Valério D. Pillar, Julieta A. Rosell, Frank M. Schurr, Serge N. Sheremetev, Ana Carolina da Silva, Ênio Sosinski, Peter M. van Bodegom, Evan Weiher, Gerhard Bönisch, Jens Kattge, Thomas W. Crowther

## Abstract

Due to massive energetic investments in woody support structures, trees are subject to unique physiological, mechanical, and ecological pressures not experienced by herbaceous plants. When considering trait relationships across the entire plant kingdom, plant trait frameworks typically must omit traits unique to large woody species, thereby limiting our understanding of how these distinct ecological pressures shape trait relationships in trees. Here, by considering 18 functional traits—reflecting leaf economics, wood structure, tree size, reproduction, and below-ground allocation—we quantify the major axes of variation governing trait expression of trees worldwide. We show that trait variation within and across angiosperms and gymnosperms is captured by two independent processes: one reflecting tree size and competition for light, the other reflecting leaf photosynthetic capacity and nutrient economies. By exploring multidimensional relationships across clusters of traits, we further identify a representative set of seven traits which captures the majority of variation in form and function in trees: maximum tree height, stem conduit diameter, specific leaf area, seed mass, bark thickness, root depth, and wood density. Collectively, this work informs future trait-based research into the functional biogeography of trees, and contributes to our fundamental understanding of the ecological and evolutionary controls on forest biodiversity and productivity worldwide.

## Introduction

Physiological and morphological traits determine the water, nutrient, and light economies of trees, directly influencing how individuals interact with each other and with the surrounding environment^1–5^. Traits that elevate performance in one habitat typically reduce performance in others, leading to selection for specific traits across environments^6^. Genetic, morphological, and biophysical constraints subsequently limit the range of traits that a species can exhibit, leading to so-called trait ‘trade-offs’ that shape species’ geographic distributions^7^, coexistence mechanisms^8,9^, and the provision of ecosystem services^10,11^. Despite a wealth of research into trait trade-offs across the plant kingdom, there remains relatively little understanding of the unique trade-offs faced by large woody species. Identifying the dominant trait trade-offs in trees is fundamental to our understanding of the functional biogeography of forests, and critical for predicting how forest diversity, composition, and function will respond to changing environmental conditions^12–15^.

Prior studies have identified a key set of traits that summarize the spectrum of form and function across herbaceous and woody plants^4,16–18^. But due to their size, longevity, ontogeny, and unique structural properties, trees have distinct characteristics and face novel abiotic stressors relative to herbaceous plants^5,19–23^. Trait analyses which include both woody and herbaceous plants are forced to omit critical aspects of tree architecture (e.g., bark properties, crown size, stem conduit diameter), which overlooks the massive energetic investments in structures that are unique to large woody species^22,24^. Understanding how tree-specific traits align with existing plant trait frameworks is key for identifying the dominant biogeographic and ecological processes governing forest structure across broad spatial scales^15^.

Here, we uses a global database^25^ of more than 350,000 trait measurements to explore relationships among 18 functional traits, reflecting leaf economics, wood structure and function, tree size and architecture, reproduction, and below-ground allocation (Fig. 1b). We asked: (1) Which dominant trade-offs and abiotic variables best capture overall variation in tree trait expression? and (2) What are the dominant multi-trait constellations that capture the breadth of form and function at the global scale? We hypothesized that traits related to canopy architecture, wood density, and tree size would emerge as key variables governing trait patterns in trees, driven by the large energetic requirements of large woody structures^26,27^. Collectively, this work identifies emergent constraints on tree functional biogeography, and sheds light on the core ecological processes shaping functional trait expression in forests worldwide.

**Figure 1.**
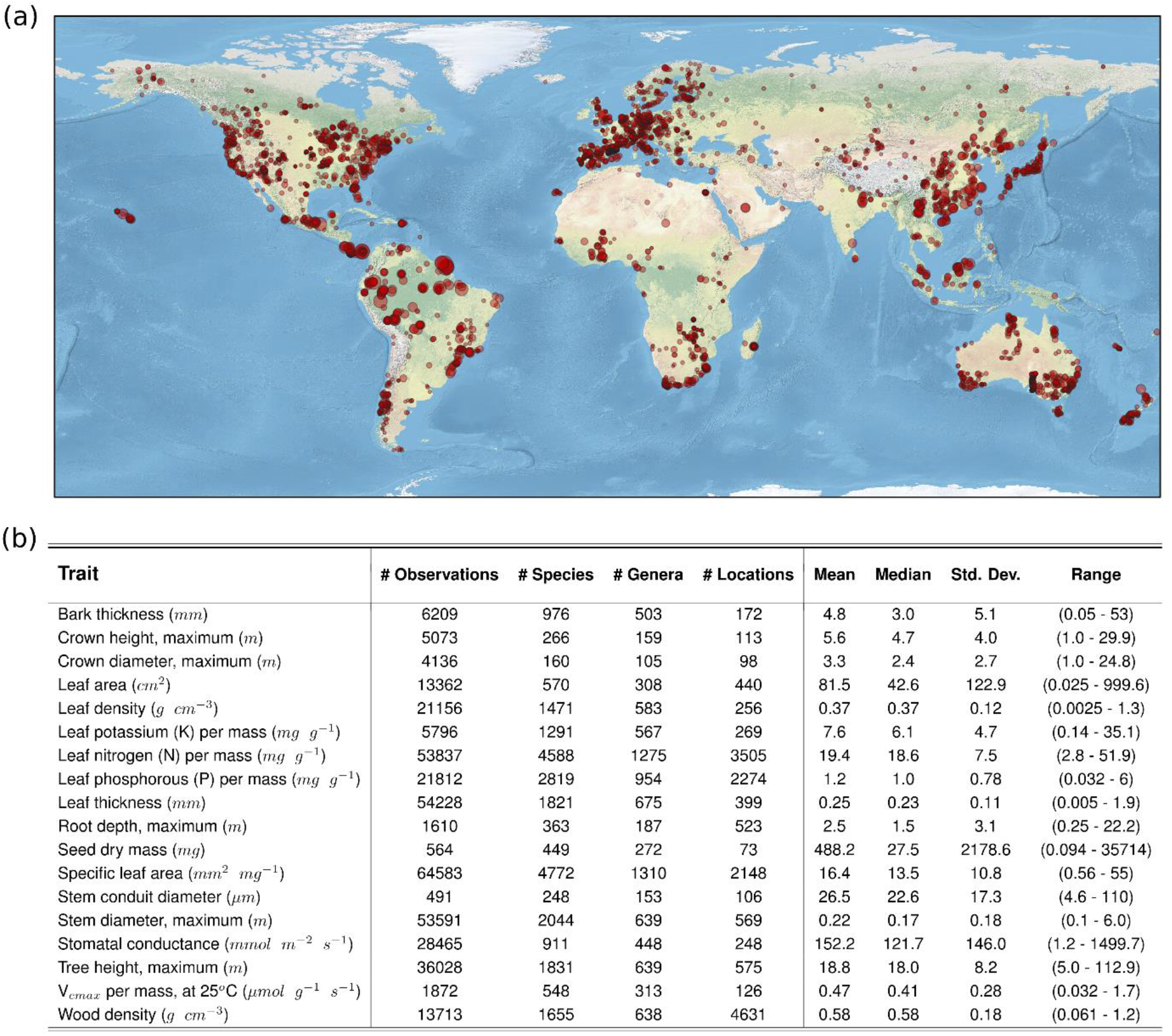
Overview of the 18 functional traits. (a) The unique geographic locations (n = 8490) where tree functional traits were recorded. The size of the circles denotes the relative number of traits measured at each location. (b) The number of unique measurements, species, locations, and genera for each of the 18 traits considered here, along with summary statistics. The analysis included 386,526 trait measurements, encompassing 6905 unique tree species and 1691 unique genera (see Table S1 for corresponding metadata).

## Results & Discussion

### TRADE-OFFS IN TREE TRAIT EXPRESSION

Our analysis included 386,526 unique trait measurements across 18 traits, encompassing 6905 tree species from 1629 genera and 203 families (Fig. 1). Traits were measured at 8490 distinct locations, including every continent except Antarctica. To explore trade-offs in functional traits at the individual level, we used random forest machine learning models to estimate trait expression for each individual as a function of its environment and phylogenetic history. Environmental predictors included a range of climate^28–31^, soil^32^, topographic^33^, and geological^34^ features. Phylogenetic history was incorporated via phylogenetic eigenvectors^35,36^ (see Methods).

Across all 18 traits, our models explained 54% of trait variation (buffered leave-one-out cross-validation, see Methods), with a relative predictive error of ± 28% (Fig. S1-S2). The inclusion of environmental variables led to substantial increases in explanatory power and accuracy, reducing the expected predictive error of the models by 10% and improving the explanatory power by 35% across all traits. Overall, environmental variables and phylogenetic information had approximately equal explanatory power (relative importance of 0.51 vs 0.49 for phylogeny vs. environment), albeit with substantial variation across traits (Fig. S3). Traits with high intraspecific variation and ontogenetic plasticity exhibited particularly strong increases in accuracy with the inclusion of environmental variables (e.g., 19% and 16% improvement for crown height and root depth, respectively). Only seed dry mass had no residual environmental signal after accounting for phylogeny (Fig. S2-S3).

Using the resulting trait models, we quantified trait trade-offs at the individual levels, accounting for both phylogenetic trait conservatism and environmental-mediated trait variation. When considering all traits simultaneously, the first two axes of the resulting principal-component analysis capture 40% of variation in overall trait expression (Figs. 2a, S4). The first trait axis correlates most strongly with leaf thickness (ρ = -0.77), specific leaf area (ρ = 0.74), and leaf nitrogen (ρ = 0.70). By capturing key aspects of the leaf economic spectrum^16^, these traits reflect various physiological controls on leaf-level resource processing, tissue turnover and photosynthetic rate^6,37,38^. Thick leaves with low SLA can help minimize desiccation, herbivory, frost damage, and nutrient limitation, but at the cost of reduced photosynthetic potential due to primary investment in structural resistance^39^. Accordingly, leaf nitrogen—a crucial component of Rubisco for photosynthesis^40,41^—trades off strongly with leaf thickness (ρ = -0.47). By reflecting an organismal-level trade-off between photosynthetic potential in optimal conditions and abiotic tolerance in suboptimal conditions, this first axis thus captures the core distinction between “acquisitive” and “conservative” functional traits which underpin fast-slow life-history strategies across the plant kingdom^6,17,18^. At one end of this spectrum are species with acquisitive traits which confer higher growth, faster nutrient cycling, and greater photosynthetic potential. At the other end are conservative traits (thick leaves) which help trees withstand a variety of stressful abiotic conditions, but which come at the cost of reduced photosynthetic capacity and growth in optimal environments.

**Figure 2.**
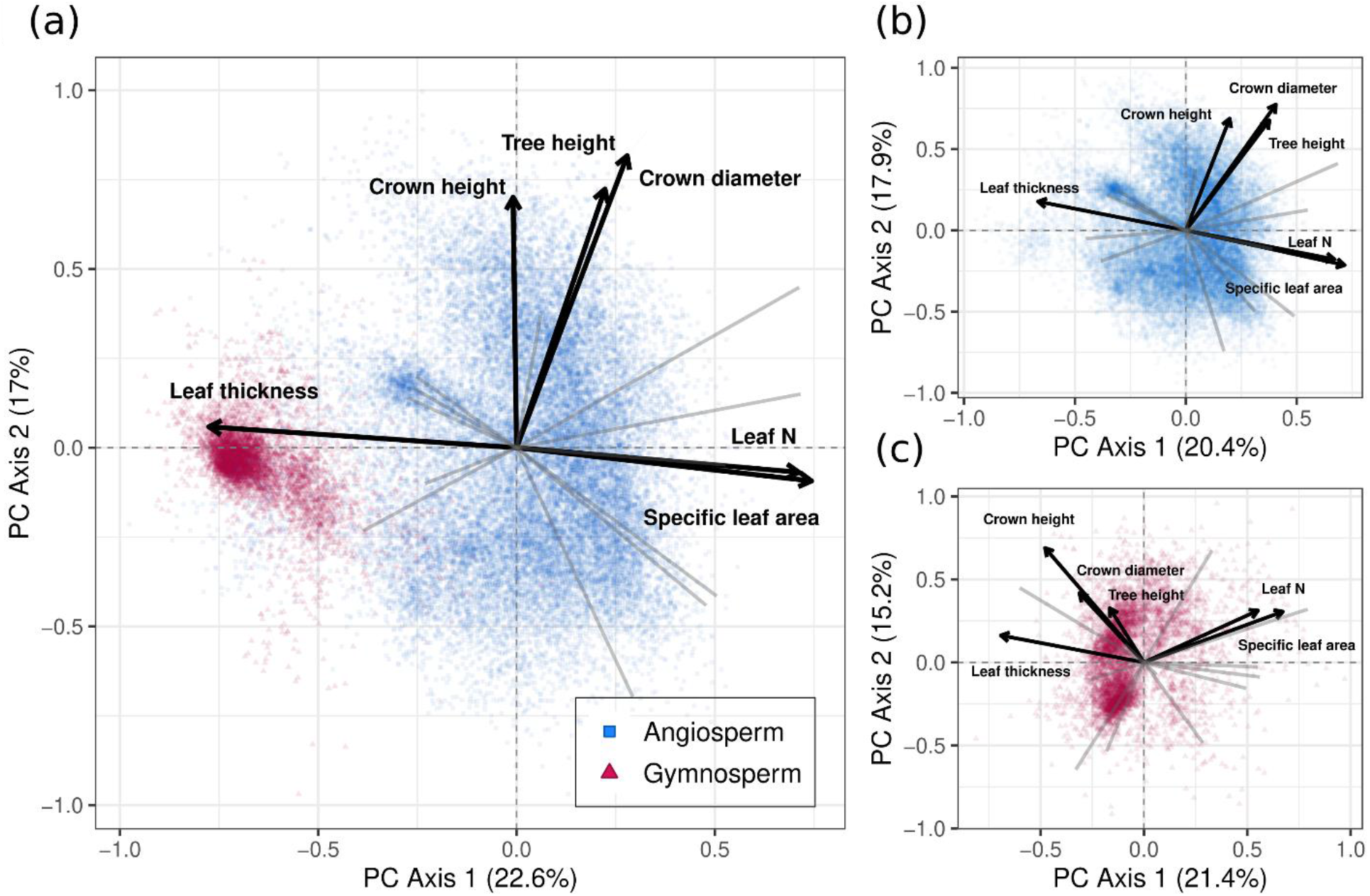
Dominant trait axes and trade-offs. Shown are the first two principal component axes capturing trait trade-offs across the 18 functional traits. (a) All tree species (n = 29,450 observations), (b) angiosperms only (n = 5457), and (c) gymnosperms only (n = 23,993). In (a) the three variables that load most strongly on each axis are show in dark black lines, with the remaining variables shown in light gray. These same six variables are shown in (b) and (c) illustrating how the same trade-offs extend to angiosperms and gymnosperms. See Supplemental Figs. S4-S6 for the full trait PCAs.

The second trait axis correlates most strongly with maximum tree height (ρ = 0.72), crown height, (ρ = 0.70), and crown diameter (ρ = 0.82), highlighting the overarching importance of competition for light and canopy position in forest ^6^ (Figs. 2a, S4). Large trees and large crowns are critical for light access and for maximizing light interception down through the canopy^42^. Nevertheless, tall trees with deep canopies also experience greater susceptibility to disturbance and mechanical damage, primarily due to wind and weight^43,44^. Because of the massive carbon and nutrient costs required to create large woody structures^26,27^, larger trees are less viable in nutrient-limited or colder climates^45^, and in exposed areas with high winds or extreme weather events^46^. This second axis thus reflects a fundamental biotic/abiotic trade-off related to overall tree size, which is largely orthogonal to leaf-level nutrient-use and photosynthetic capacity.

Despite substantial differences in wood and leaf structures between angiosperms and gymnosperms (e.g., vessels vs. tracheids), the two main trade-offs hold within, as well as across, clades (Fig. 2b-c, S5-S6). Gymnosperms, however, exhibit less orthogonality between these axes, in part due to less variation in leaf thickness and a stronger subsequent correlation between leaf thickness and tree size (ρ = 0.39 vs. 0.04 for gymnosperms vs angiosperms). Nevertheless, despite differences in physiology and morphology, gymnosperms and angiosperms are subject to the same physical, mechanical, and chemical processes that determine the ability to withstand various biotic and abiotic pressures^7,47,48^. Our results show that these processes translate into similar fundamental constraints on trait expression across clades.

Collectively, the two primary trait axes thus reflect two different aspects of the dominance-tolerance trade-off: (1) the ability to maximize leaf photosynthetic activity, at the cost of increase risk of leaf desiccation, and (2) the ability to compete for space and maximize light interception via tree size, at the cost of increased susceptibility to mechanical damage. Notably, these two trade-offs closely mirror those seen when considering herbaceous species alongside woody species^4^, though we observe stronger orthogonality between leaf function and plant size is found when considering the whole plant kingdom. Thus, rather than fundamentally reshape the dominant trade-offs, the inclusion of tree-specific traits shows that these two ecological constraints systematically affect all aspects of tree form and function, illustrating the universality of these two ecological trade-offs across the plant kingdom.

### Environmental predictors of trait trade-offs

To examine how environmental variation shapes trait expression across the globe, we quantified the relationships between environmental conditions and the dominant trait trade-offs.

In line with previous analysis^49^, temperature variables were the strongest univariate drivers of trait trade-offs (Figs. 3, S7). The first PC axis (leaf thickness) correlates most strongly with annual mean temperature (ρ = 0.26, Fig. 3a), reflecting that leaves face increased frost risk and reduced photosynthetic potential in colder conditions. Thus, selection should favor thick leaves with low SLA over thin leaves with high SLA and high nutrient-use^37^. However, the univariate environmental signal is relatively weak (Fig. 3c), highlighting that the first PC axis captures more complex relationships among leaf-level economies and environmental conditions (e.g. between angiosperms vs. gymnosperms).

**Figure 3.**
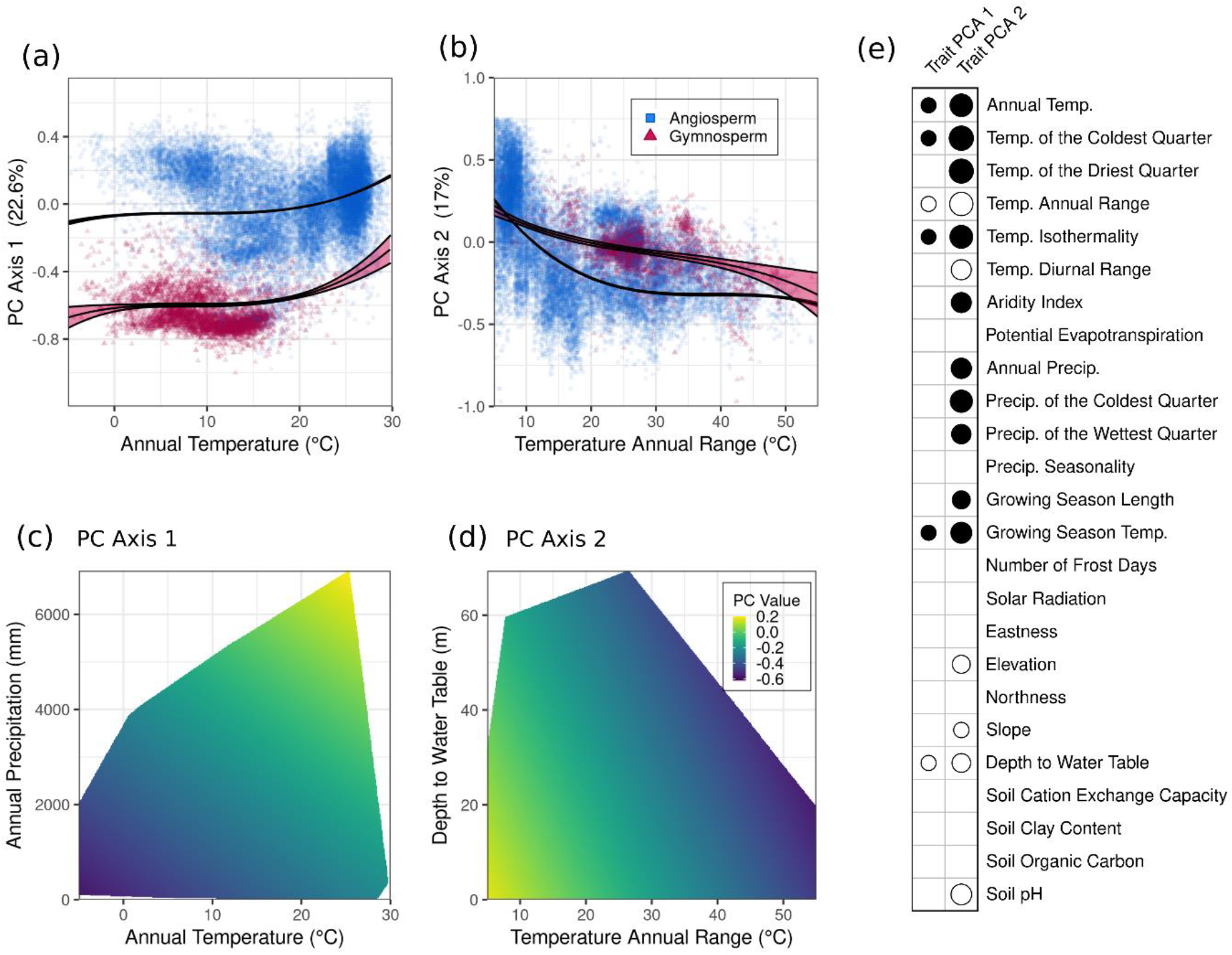
The relationship between environmental variables and trait trade-offs. (a-b) The strongest univariate predictor of each trait trade-off. (c-d) The 2-dimensional surfaces showing the strongest bivariate predictors of each trait trade-off. The surfaces are constrained to the convex hull of the observed variable combinations in the dataset. Note that the x-axes in *c-d* align with those above in *a-b*. (e) Correlations between the first two trait trade-offs and environmental variables. The size of the circle denotes the relative strength of the correlation, with solid circles denoting positive correlations and open circles denoting negative correlations. For clarity, only trait correlations with |ρ| > 0.2 are shown. See Fig. S7 for the full set of correlations.

The second PC axis (tree size) correlates most strongly with temperature annual range (Fig. 3b). Crown diameter, leaf area, and tree height, in particular, all exhibit strong negative correlations with temperature annual range (ρ = -0.65, -0.56 and -0.54, respectively, Fig. 3c). At the global scale, high temperature variation is inversely correlated with annual temperature (ρ = -0.79), such that larger structural components are favored in areas with consistently warm temperatures— primarily tropical regions near the equator, and coastal regions throughout the Americas, Australia, and southern Africa. Trees in such environments are more likely to experience strong biotic interactions, which should increase evolutionary and ecological selection pressures over time^50,51^, favoring species with high competitive ability and efficient light acquisition strategies.

Despite the primary importance of temperature governing tree trait trade-offs, precipitation and temperature regimes are highly correlated, and the main climate stressors to trees arise via interactions between temperature and water availability (e.g., xylem cavitation and embolism, fire regimes, and leaf desiccation). Indeed, when exploring the bivariate drivers of trait expression, precipitation variables emerge as the strongest secondary drivers of each trait trade-off. The first PC axis exhibits the highest values at high temperatures in combination with high precipitation (Fig. 3c), reflecting broad-scale differences in habitat requirements across angiosperms and gymnosperms^52,53^. For the second PC axis, higher values are observed among trees in regions with low temperature variation in tandem with sufficient ground water access (Fig. 3d), demonstrating that soil hydrology and precipitation place key limitations on tree size at the global scale^54^.

### Trait constellations at the global scale

Although exploration of trait PC axes sheds light on the dominant physiological trade-offs structuring tree traits, these first two trait axes account for less than half of overall trait variation (Fig. 2a). To better explore the multidimensional nature of these trade-offs, we subsequently identified groups of traits that form tightly coupled clusters and which reflect distinct aspects of tree function.

Our results show that these 18 traits can be grouped into seven trait constellations, each of which reflects a unique aspect of tree growth, physiology, or ecology (Fig. 4). The largest trait constellation (Fig. 4, pink cluster) loads most heavily on the first trait trade-off (Fig. 4b), capturing various aspects of the leaf economic spectrum. Relationships among specific leaf area, leaf V_cmax_ (the maximum rate of carboxylation), leaf thickness, and leaf nutrient concentrations per mass (N, P, K) are well established^4,6,41,55^. The fact that nearly all leaf traits fall in this cluster (with the exception of leaf area and density) supports the inference that leaf economics represent a unique aspect of tree function that is largely independent of plant size ^4^.

**Figure 4.**
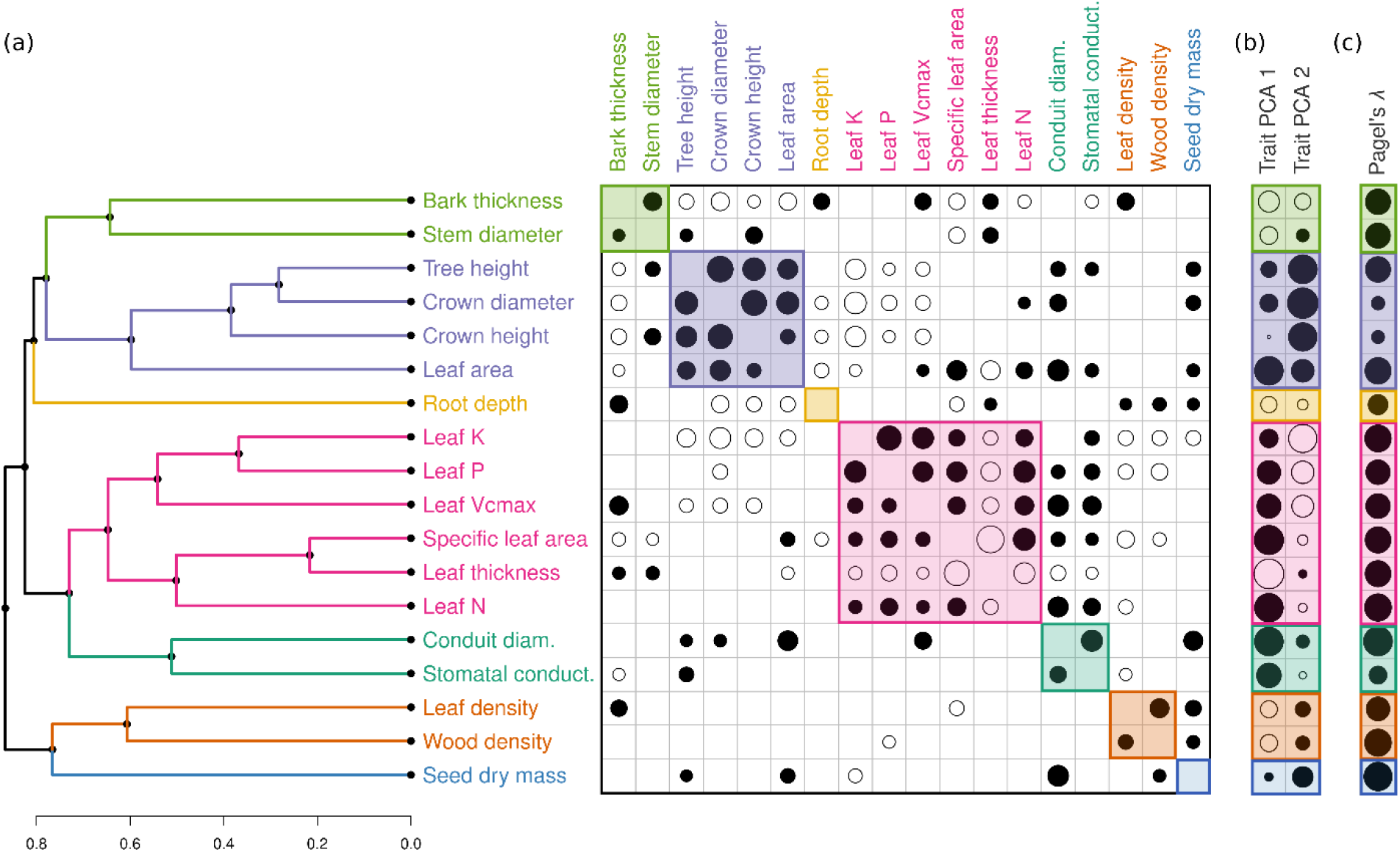
Trait correlations and functional constellations. (a) Trait clusters with high average intra-group correlation. The upper triangle shows species-weighted correlations incorporating intraspecific variation, and the lower triangle gives the corresponding correlations among species-level phylogenetic independent contrasts. The size of the circle denotes the relative strength of the correlation, with solid circles denoting positive correlations and open circles denoting negative correlations. For clarity, only trait correlations with |ρ| > 0.2 are shown. (b) Correlations between each trait and each of the first two principal component axes, illustrating which functional trait clusters align most strongly with the dominant axes of trait variation. (c) The species-level phylogenetic signal of each trait, calculated using only the empirical trait measurements.

The second-largest trait constellation (Fig. 4, purple cluster) includes tree-size traits which closely align with the second trade-off, along with the addition of leaf area. As with tree height and canopy size, leaf area directly affects a tree’s ability to intercept light down through its canopy^42^. Leaf area thus serves as an intermediary between the two primary trait axes: it is intrinsically correlated with SLA but exhibits relatively weak correlations with per-mass leaf nutrients. Although the largest discrepancies in leaf area are observed between needle-leaf gymnosperms and broadleaf angiosperms, these trends hold within clades as well, with leaf area among gymnosperms likewise correlating positively with both tree height (ρ = 0.20) and crown diameter (ρ = 0.48). This cluster thus highlights organismal-level coordination of light interception that integrates tree size, architecture, and leaf shape.

Intermediate to these two largest clusters are three constellations each containing two traits: (1) stem conduit diameter and stomatal conductance (Fig. 4, dark green), capturing organismal-level integration of water transport at the cost of increased desiccation and cavitation risks; (2) stem diameter and bark thickness (Fig. 4, light green), primarily demonstrating intrinsic size-based relationships between stem parts^56,57^, which are secondarily related to aridity and fire frequency in some environments^57^; and (3) wood density and leaf density (Fig. 4, orange), indicative of slow/fast life-history strategies, where denser plant parts reduce growth rate and water transport^5,17^ but protect against pest damage, desiccation, and mechanical breakage^5,27,39^. Collectively, these two-trait clusters each demonstrate unique and complementary trade-offs that insulate trees against various disturbances and extreme weather events, but at the cost of reduced growth, competitive ability, and productivity under optimal conditions (see Supplemental Discussion).

Lastly, two traits each comprise their own unique cluster: root depth and seed dry mass (Fig. 3, yellow and blue). Root growth is subject to a range of belowground processes (e.g., root herbivory, depth to bedrock) that can promote a disconnect between aboveground climate conditions and belowground traits^54,58,59^. Root depth accordingly has a relatively weak phylogenetic signal (λ = 0.44) but a strong environmental signal (Figs. 4, S1-S2), reflecting distinct belowground constraints on trait expression. In contrast, seed dry mass exhibits the strongest phylogenetic signal (λ = 0.98, Fig. 4c) and weakest environmental signal of any trait (Figs. S1-S2). Reproductive traits are subject to unique evolutionary pressures^60^, indicative of different seed dispersal vectors (wind, water, animals) and various ecological stressors that uniquely affect seed viability and germination^60^. The emergence of root depth and seed mass as solo functional clusters thus supports previous inference that belowground traits and reproductive traits reflect distinct aspects of tree form and function not captured by leaf or wood trait spectrums.

Collectively, these trait constellations shed light on organismal-level trait coordination and broad-scale differences in trait expression across species and clades. A key challenge in identifying global patterns in trait trade-offs is the relatively sparse trait coverage at the individual level, with only a handful of traits typically measured on any single tree. This limitation is partly due to the enormous range of putative traits that can be measured on trees^6,25^. Here, by using phylogenetic and environmental information to estimate trait expression at the individual level, our approach helps to overcome some of these limitations, enabling us to explore organismal level trade-offs across thousands of species. The benefit of this approach we can include vastly broader phylogenetic, geographic, and trait coverage than would otherwise be possible. Future work, however, should focus on improving the precision of these global trait frameworks by measuring complete sets of traits on individual trees.

To help address these challenges, the seven described trait constellations (Fig. 4) can be used as a starting point for research into organismal-level trait expression in trees. Although the exact subset of traits used in a given study should depend on the intended scope and application, we advise selecting traits which exhibit strong phylogenetic signal and/or low environmental-mediated variation, and ideally have low cross-correlations with other traits in other clusters. These criteria help to ensure that the selected traits can be robustly measured and that they reflect well-defined ecological processes. In line with this, we suggest a baseline set of seven traits selected from these trait constellations: bark thickness, maximum tree height, root depth, specific leaf area, stem conduit diameter, wood density, and seed dry mass. These seven traits represent complementary ecological and evolutionary processes, capturing differences in competitive ability, growth rate, abiotic stress tolerance, reproduction, wood and leaf properties, and above-vs. belowground allocation. Moreover, these traits are relatively well represented in many trait databases and have well-defined definitions and measurement protocols^25,61,62^, thus forming a baseline set of reference traits for expanding our understanding of tree functional trait expression.

## Conclusions

Collectively, our analysis reveals key trade-offs and trait constellations governing tree form and function worldwide. We show that tree functional traits predominantly reflect two major functional trade-offs: one representing leaf-level nutrient-use and photosynthesis, and the other representing competition for light via tree and crown size. Mirroring patterns seen across the entire plant kingdom, these trade-offs capture an ecological gradient from conservative growth strategies under suboptimal environments (cold, dry, frequent disturbances), to acquisitive strategies associated with light competition in high-resource environments (consistently high temperature and water availability). By incorporating traits unique to large woody species, we further identify a unique set of functional constellations and representative traits that reflect the breadth of tree form and function. In doing so, these results elucidate key constraints on functional trait relationships in trees, contributing to our fundamental understanding of the controls on the function, distribution, and composition of forest communities. By identifying a core set of traits that reflect the broad variety of ecological life-history strategies in trees, this work can inform future trait-based research into the functional biogeography of the global forest system.

## Data and code availability

The data and code for replicating the central findings will be made available in a dedicated GitHub repository upon publication.

## Author contributions

DSM conceived of this study and analyzed the data, with assistance from LB, CMZ, CA, JvdH, HM, LM, GRS and TWC. Data were contributed by IA, EB, CCFB, JC, BELC, ASD, AG-M, PH, CHL, ASK, ÜN, VDP, JAR, FMS, SS, ACdS, ÊS, PMvB, EW, GB, and JK, who also provided suggestions and feedback on the analyses and interpretations. All authors contributed to the writing and revising of the manuscript.

## Acknowledgements

This work was primarily funded by the Swiss National Science Foundation, Ambizione grant #PZ00P3_193612 to DS Maynard. Additional support was provided by D.O.B. Ecology to TW Crowther; SNSF Ambizione grant #PZ00P3_193646 to CM Zohner; SNSF Ambizione grant # PZ00P3_179900 to C Averill; U.S. National Science Foundation Graduate Research Fellowship and Embassy of Switzerland in the USA ThinkSwiss Research Fellowship to GR Smith; and UNAM-PAPIIT grant #IN210220 to JA Rosell. The authors thank Owen Atkin and Yongfu Chai for additional data, as well as the TRY database managers and numerous researchers who contributed open-access data.

## Materials & Methods

### Trait information

Trait data were obtained from the TRY plant trait database^1^ in April 2020. Data were cleaned by converting all traits to standardized units and by matching species names to The Plant List (TPL) database v1.1 (http://www.theplantlist.org, accessed June 2020) using the *Taxonstand* package in R v3.6.0^2^. Synonyms were replaced with accepted names, when available. The phylogenetic tree was taken from the seed plant phylogeny of Smith & Brown (2018), and species names were likewise cleaned and harmonized using the TPL database. To limit our analysis to tree functional traits only, we used the BGCI GlobalTreeSearch database v1.3^4,5^, containing a comprehensive list of ca. 60,000 tree species compiled and harmonized from across 500 sources. The BGCI database uses TPL for much of its taxonomic identification, but to ensure consistency among all sources we used the same name harmonization pipeline as with TRY and the seed plant phylogeny. We constrained the set of traits and the phylogenetic tree to those species that could be matched to the BGCI database (n = 54,153 species matched), and we likewise trimmed the phylogenetic tree to the set of species matched in TRY. We further excluded any trait observations that did not have corresponding geographic coordinates.

Traits were selected based on data availability, phylogenetic and spatial coverage, and importance for tree growth, survival, and competition. We prioritized traits that are commonly used in other leaf and wood economic spectrums^6^, thus focusing on leaf traits measured per unit mass rather than unit area, where possible. Secondly, we prioritized traits which are important indicators of tree growth and structure, such as stem conduit diameter and crown dimensions. Thirdly, we selected traits which capture important ecological processes and life-history differences among trees (e.g., root depth). This process resulted in 30 putative traits for analysis (Table S1, Supplemental Data References). Of these, we omitted traits with low sample sizes (either overall, or at the species, genus, or geographic level), those that are redundant or intrinsically correlated with other traits, and those that are known to be highly sensitive to trait assay conditions but were not clearly standardized. This pruning resulted in a set of 18 focal traits for use in the final analysis, with the 12 omitted traits used only to improve predicted power via trait covariation (see model process, below). Trait values were converted to common units where necessary (e.g., mm to cm). For each trait, we selected sub-categories (as given by TRY) that denoted comparable measurements and reflected uniform assay conditions (e.g., V_cmax_ measured at 25°C) (Table S1).

### Model details

In order to consider trait trade-offs at the organismal level which accounted for intraspecific variation, we used machine learning models to estimate all 18 trait values for each individual tree in each location (Fig. 1a). We modeled trait expression as a function of both environmental and phylogenetic information so as to estimate traits with weak phylogenetic signals but strong abiotic filtering, and to incorporate intraspecific variation and ontogenetic plasticity into our analysis. Specifically, we used random forest (RF) models to estimate trait values for each observation using the *ranger* package in R^7^. Initial exploration showed negligible effects of model tuning, such that the default hyper-parameters were used to prevent overfitting. In order to minimize the influence of data-recording errors or unit mismatches in the dataset, trait values which occurred outside of the bulk of the trait distribution were investigated as outliers. Those which could not be externally verified and which were biologically unreasonable were removed (e.g., stem diameters >15 m). When modeling tree height, canopy size, and root depth, we only considered observations with height >5 m, diameter >10 cm, root depth >25 cm, and canopy dimensions >1 m high and wide^8^, thereby ensuring that our analysis focused on adult trees rather than saplings or woody shrubs. We subsequently implemented quantile random forest^9,10^ to estimate the upper 90^th^ percentile trait value for maximum stem diameter, canopy dimensions, and root depth. In all other cases the imputed traits represent the mean predicted value across the random forest.

Environmental covariates used in the models included 50 variables encompassing range of climate^11–13^, soil^14^, topographic^15^, and geological^16^ variables (Table S2). We omitted variables that directly measure plant community composition or biotic factors (e.g., NDVI or % forest cover) so as to ensure the resulting geographic layers solely encompassed abiotic factors. Layers were sampled from a previously prepared global composite (see van den Hoogen et al. 2019 for details). Briefly, all covariate map layers were resampled and reprojected to a unified pixel grid in EPSG:4326 (WGS84) at 30 arcsec resolution (approximately 1 km^2^ at the equator). Layers with a higher original pixel resolution were downsampled using a mean aggregation method; layers with a lower original resolution were resampled using simple upsampling (that is, without interpolation) to align with the higher resolution grid. The set of environmental covariates for each trait measurement was obtained by sampling this composite image at each unique latitude and longitude value given in the TRY database.

Phylogenetic information was incorporated in the form of phylogenetic eigenvectors^18–21^. We first calculated the pairwise cophenetic phylogenetic distance matrix across all 54,153 tree species that could be matched to both the BGCI tree list and the plant phylogeny. This matrix was then double-centered by rows and columns^21,22^, and the first 50 orthogonal eigenvectors were extracted from this matrix for use as continuous predictors in the random forest models. The choice of 50 eigenvectors (out of 54,153 possible) was in line with previous analyses to prevent over-fitting and to ensure the model was identifiable^22^. This also resulted in the same number of environmental and phylogenetic predictor variables.

To leverage trait covariation among the disparate observations, we used a two-step algorithm to improve predictive power and imputation accuracy^20,23^. First, following standard approaches^6,24–26^, trait values were log-transformed, allowing for comparisons across trait distributions which are highly right-skewed and vary by several orders of magnitude^26^ (Fig. 1). Using the general approach of Stekhoven & Bühlmann (2012), we next implemented a random forest on all traits for all observations. We then used these initial models to predict the full set of trait values for each observation (including the 12 ancillary traits not included in the focal analysis, Table S1). We then refit the random forest models for each trait, using the full set of predicted traits (apart from the focal traits) as covariates. For the final analysis, observed traits were used in place of imputed traits, when available, with the exception of maximum tree height, stem diameter, root depth, and crown size, where the upper 90^th^ percentile trait values were used. Variable importance in the random forest models was calculated using the “permutation” metric, reflecting the variance in responses across predictors^7^.

### Model performance

Model performance was quantified using buffered leave-one-out cross-validation^27^. To avoid overfitting, we followed the approach recommended in Roberts *et al*. (2017) and fit a simple linear model to the data, where trait expression was modeled as a linear function of phylogenetic and environmental covariates. We then assessed spatial autocorrelation of the residuals using Moran’s I plots using the *ncf* package in R, which displays the value of spatial autocorrelation (ranging from -1 to 1) as a function of distance^28^. We likewise assessed residual phylogenetic autocorrelation across taxonomic ranks (genus, family, order, group), using the the *ape* package in R. In general, spatial autocorrelation was low (I<0.10) (Fig. S8), with the exception of leaf phosphorous, which exhibited slight autocorrelation up to ∼250 km. Residual phylogenetic correlation was likewise low, and generally only observable at the genus level, apart from crown size and conduit diameter, which exhibited residual autocorrelation up to the family level (Fig. S9). Thus, to be conservative, for all traits except crown size and conduit diameter, we used a genus-level spatial buffer of 250 km to exclude test/training data; and for crown size and conduit diameter we used a family-level buffer at 250 km. To implement the cross-validation accuracy assessment, we first randomly selected a focal species, with the out-of-fit test data containing all observations for that species for the focal trait. To construct the corresponding training data, we excluded all observation of the same genus (or family) that fell within a 250km spatial buffer of any of the training points for that species. The random forest models were then fit using the buffered training data, and used to predict the average trait value for the omitted species^27^. This procedure was repeated for each unique species for each trait, up to 1000 times, with a randomly sampled focal species selected at each iteration.

Predictive accuracy was assessed in two ways. First, following the recommendation of Li (2017), we calculated the cross-validated coefficient of determination relative to the 1:1 line (termed “VEcv”, Li 2017), which provides a normalized version of the mean-squared-error (MSE) that allows for comparisons across data types and units. Specifically, this value is calculated as: 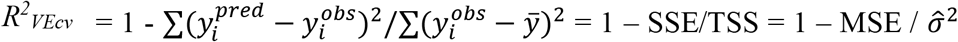, where the summation is taken across the species, and the predicted values are estimated out-of-fit using the buffered cross-validation procedure outlined above. Importantly, this metric is not the same as a regression-based goodness-of-fit, as it is calculated by direct comparison of observed vs. predicted values^29^. Second, we also report the median absolute percentage error (MdAPE), which gives a more interpretable estimate of the expected error a given prediction, calculated as 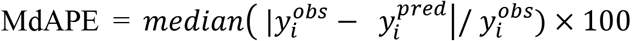. Although the models were fit using log-transformed data, accuracy was assessed on the non-logged values in their original units.

### Principal component analysis

Species-weighted principal component analysis (PCA) was conducted on the full set of imputed traits using the *aroma*.*light* package in R. The weights were set to be inversely proportional to the number of observations for each species, which allowed us to incorporate intraspecific variation while also ensuring that each species had the same overall contribution to global trade-offs. Representative vectors for each axis were identified by selecting those that correlated most uniquely on each of the first two principle component axes.

### Abiotic relationships

To identify univariate and bivariate relationships among trait trade-offs and environmental conditions, we first identified the environmental variable that correlated most strongly with each of the two PC axes (Fig. 3a-b, Fig. S7). We used Spearman rank correlations to allow for nonlinear relationships among traits and environmental conditions. To visualize these correlations, we separately fit third-order monotonic regression polynomials for angiosperms and gymnosperms, and obtained 95% bootstrap confidence intervals by randomly sampling one observation for each species per iteration, repeated 500 times. To explore the bivariate predictors of trait trade-offs, we then fit a series of simple pairwise linear regression models to identify which additional environmental variable led to the highest subsequent increase in explanatory power for each trait (measured via adjusted R^2^). To avoid spurious relationships due to the large number of pairwise combinations (50 choose 2), we only considered a subset of representative environmental traits, identified via cluster analysis: annual precipitation, annual temperature, temperature annual range, precipitation seasonality, aridity index, growing season mean temperature, growing season length, permafrost extent, soil water-holding capacity, soil cation-exchange capacity, soil pH, topographic northness, topographic eastness, and depth to water table. To visualize the resulting patterns (Fig. 3c-d), we plotted the smooth regression surfaces across the full range of environmental conditions, restricted to the convex hull of the observed variable combinations in the dataset.

### Hierarchical cluster analysis

Trait cluster analysis was conducted using hierarchical clustering on the species-level correlation matrix. First, we calculated species-weighted rank correlations between pairs of traits using the *wCorr* package in R, which again allowed us to incorporate intraspecific trait variation while ensuring each species contributed equal weight. The optimal number of clusters was identified using the silhouette method in the *dendextend* package in R, and the dendrogram was subsequently cut into clusters based on groups of traits which exhibited consistently high average intra-group correlation. As an alternate measure of trait correlation which accounts for phylogenetic relatedness, we calculated phylogenetic independent contrasts^30^ on species-level average trait values using the *ape* package. The corresponding correlations among these contrasts are shown in the bottom triangle of the correlation matrix in Fig. 4. Species-level phylogenetic conservatism was calculated via Pagel’s λ, using only the empirically measured values in the TRY dataset.

All analyses were conducted in R v. 3.6.0, with the exception the phylogenetic eigenvector calculations, which were obtained using the *Arpack* package in Julia v. 1.6.2.

