## Supplemental Material for "Global trade-offs in tree functional traits"

Daniel S. Maynard *et al.*

Institute of Integrative Biology, ETH Zürich, 8092 Zürich, Switzerland

**CONTENTS:**

- Supplemental Discussion
- Table S1-S2
- Figures S1-S9
- Supplemental References
- Supplemental Data References

### Supplemental Discussion

*(Here we provide additional context for the three 2-trait clusters in Figure 4, main text)*

Intermediate to the two largest clusters in Fig. 4 (main text) is a constellation containing stem conduit diameter and stomatal conductance, demonstrating leaf/wood water regulation (Fig. 4, yellow). This constellation loads most strongly on PC 1, as moisture regulation, nutrient-use, and photosynthesis are closely interrelated. Nevertheless, conduit diameter, in particular, correlates moderately with traits indicative of light interception, notably leaf area ($\rho = 0.48$) and tree height ($\rho = 0.25$). Although the largest differences in stem conduit size are observed between angiosperms and gymnosperms, wider conduits confer greater conducting efficiency regardless of architecture[1]. Indeed, these patterns hold within clades as well, particularly when comparing conduit diameter to leaf area ($\rho = 0.43$ vs 0.38 for angiosperms vs. gymnosperms). The associations between water regulation and light interception highlights that leaf area and tree height induce important physiological and mechanical demands on organism-level water availability, which prevent against cavitation and desiccation.

Tree diameter and bark thickness also emerge as a distinct two-trait cluster. Recent research has shown that this relationship is predominantly an intrinsic byproduct of overall tree size[2], partly because, in many species, bark thickness increases monotonically as trees age due to continued fissuring and production of new bark[3]. From an ecological perspective, thick bark can be critical for defense against fire and pest damage (mainly a thick outer bark region), and for storage (mainly a thick inner bark region)[4–7]. Moreover, because older individuals are more likely to have been exposed to multiple disturbances across their lifetime, large-diameter trees in older forests can exhibit survivorship bias towards thick-bark individuals which were able to withstand historical stressors[3,8]. However, such relationships are strongly ecosystem-dependent, leading to weak overall relationships between climate, fire regimes, and bark thickness at the global scale, with stem diameter emerging as the strongest single predictor[2].

The final two-trait cluster is comprised of wood density and leaf density. Both traits are key indicators of “slow” life-history strategies in trees, correlating negatively with growth rate and water transport, but positively with abiotic stress tolerance and resilience to disturbance[9]. Thick leaves and thick wood each protect against herbivory and pests while protecting against desiccation risk and mechanical damage[10–13]. Wood density has been identified as a particularly important multi-functional indicator of tree form and function, reflecting various aspects of tree hydraulics, pest resistance, decay rate, structural stability, tree size, growth rate, and tree mortality[11,14–17]. The production of dense wood and leaves is, however, more energetically costly, limiting growth rate but increasing life span[14,18]. The fact that wood and leaf density emerge as an independent trait constellation reinforces previous inference that these traits are aligned at one end of the slow-fast spectrum, and uniquely integrate multiple physiological and ecological pressures[9,14,18].

**Table S1.** The 30 functional traits and corresponding TRY trait IDs and sub-trait IDs. Of these 30 putative traits, the 18 traits used in final the analysis (indicated with *) were selected based on uniqueness, consistency of assay conditions, taxonomic coverage, geographic coverage, and overall sample size. Note that several traits (e.g., Trait 4, wood density) have multiple corresponding sub-trait IDS.

| TRY Trait ID | Sub-trait Data IDs | Trait Name |
|---:|---:|---|
| 4* | 4<br>1629<br>1739<br>2568 | Stem specific density (SSD) or wood density |
| 6* | 7 | Root rooting depth |
| 9 | 10 | Root/shoot ratio |
| 14* | 15 | Leaf nitrogen (N) content per leaf dry mass |
| 15* | 16 | Leaf phosphorus (P) content per leaf dry mass |
| 21* | 24 | Stem diameter |
| 24* | 28 | Bark thickness |
| 26* | 30 | Seed dry mass |
| 41 | 46 | Leaf respiration rate in the dark per leaf dry mass |
| 44* | 49 | Leaf potassium (K) content per leaf dry mass |
| 45* | 50 | Stomatal conductance per leaf area |
| 46* | 53 | Leaf thickness |
| 48* | 55 | Leaf density (leaf tissue density, leaf dry mass per leaf volume) |
| 56 | 100 | Leaf nitrogen/phosphorus (N/P) ratio |
| 80 | 272 | Root nitrogen (N) content per root dry mass |
| 144 | 446 | Leaf length |
| 145 | 447 | Leaf width |
| 146 | 455 | Leaf carbon/nitrogen (C/N) ratio |
| 185* | 549<br>2382 | Leaf photosynthesis carboxylation capacity (Vcmax) per leaf dry mass |
| 270 | 664 | Leaf photosynthesis electron transport capacity (Jmax) per leaf dry mass |
| 281* | 713 | Stem conduit diameter (vessels, tracheids) |
| 324* | 818 | Crown (canopy) length: diameter along the longest axis |
| 413 | 996 | Leaf chlorophyll content per leaf area |
| 773* | 1695 | Crown (canopy) height (base to top) |
| 1055 | 1950 | Root carbon/nitrogen (C/N) ratio |
| 1229 | 2657 | Wood nitrogen (N) content per wood dry mass |
| 3106* | 19<br>448<br>504 | Plant height vegetative |
| 3110* | 6577 | Leaf area (in case of compound leaves: leaf, petiole included) |
| 3117* | 6584<br>6598 | Leaf area per leaf dry mass (specific leaf area, SLA or 1/LMA) |
| 3120 | 2261 | Leaf water content per leaf dry mass (not saturated) |

**Table S2**. The environmental covariates used as predictors in the random forest models.

| Variable | Source | Type | Units | Resolution |
|---|---|---|---|---|
| Annual Temp. | 1 | Climatic | ºC | 30 arcsec (≈900m at equator) |
| Temp. of the Coldest Quarter | 1 | Climatic | ºC | 30 arcsec (≈900m at equator) |
| Temp. of the Driest Quarter | 1 | Climatic | ºC | 30 arcsec (≈900m at equator) |
| Temp. of the Warmest Quarter | 1 | Climatic | ºC | 30 arcsec (≈900m at equator) |
| Temp. of the Wettest Quarter | 1 | Climatic | ºC | 30 arcsec (≈900m at equator) |
| Temp. Annual Range | 1 | Climatic | ºC | 30 arcsec (≈900m at equator) |
| Temp. Seasonality | 1 | Climatic | ºC | 30 arcsec (≈900m at equator) |
| Temp. Isothermality | 1 | Climatic | Unitless | 30 arcsec (≈900m at equator) |
| Temp. Diurnal Range | 1 | Climatic | ºC | 30 arcsec (≈900m at equator) |
| Aridity Index | 2 | Climatic | AI Value | ≈1km |
| Potential Evapotranspiration | 2 | Climatic | PET Value (mm) | ≈1km |
| Annual Precip. | 1 | Climatic | mm | 30 arcsec (≈900m at equator) |
| Precip. of the Coldest Quarter | 1 | Climatic | mm | 30 arcsec (≈900m at equator) |
| Precip. of the Driest Quarter | 1 | Climatic | mm | 30 arcsec (≈900m at equator) |
| Precip. of the Warmest Quarter | 1 | Climatic | mm | 30 arcsec (≈900m at equator) |
| Precip. of the Wettest Quarter | 1 | Climatic | mm | 30 arcsec (≈900m at equator) |
| Precip. Seasonality | 1 | Climatic | mm | 30 arcsec (≈900m at equator) |
| Potential Evapotranspiration (std. dev.) | 1 | Climatic | PET Value (mm) | 30 arcsec (≈900m at equator) |
| Relative Humidity | 1 | Climatic | % * 100 | 30 arcsec (≈900m at equator) |
| Relative Humidity (std. Dev.) | 1 | Climatic | % * 100 | 30 arcsec (≈900m at equator) |
| Growing Season Length | 1 | Climatic | number of days | 30 arcsec (≈900m at equator) |
| Growing Season Length (std. dev.) | 1 | Climatic | number of days | 30 arcsec (≈900m at equator) |
| Growing Season Temp. | 1 | Climatic | ºC | 30 arcsec (≈900m at equator) |
| Growing Season Temp. (std. dev.) | 1 | Climatic | ºC | 30 arcsec (≈900m at equator) |
| Number of Frost Days | 1 | Climatic | Number of days | 30 arcsec (≈900m at equator) |
| Number of Snow Days | 1 | Climatic | number of days | 30 arcsec (≈900m at equator) |
| Solar Radiation | 1 | Climatic | kJ m-2 | 30 arcsec (≈900m at equator) |
| Solar Radiation (std. dev.) | 1 | Climatic | kJ m-2 | 30 arcsec (≈900m at equator) |
| Cloud Cover | 3 | Climatic | % cloudy days | 30 arcsec (≈900m at equator) |
| Cloud Cover (std. dev.) | 3 | Climatic | % cloudy days | 30 arcsec (≈900m at equator) |
| Burnt Areas (probability) | 4 | Climatic | Proportion of burned areas | ≈500m |
| Snow (probability) | 4 | Climatic | Proportion of snow occurrence | ≈500m |
| Permafrost Extent | 5 | Climatic | Unitless | 30 arcsec (≈900m at equator) |
| Depth to Water Table | 6 | Geological | m below land surface | 30 arcsec (≈900m at equator) |
| Depth to Bedrock | 7 | Geological | cm (up to 200) | ≈250m |
| Soil Bulk Density | 7 | Soil | kg / cubic–meter | ≈250m |
| Soil Cation Exchange Capacity | 7 | Soil | cmolc/kg | ≈250m |
| Soil Clay Content | 7 | Soil | mass fraction in % | ≈250m |
| Soil Coarse Fragments | 7 | Soil | % | ≈250m |
| Soil Water Capacity | 7 | Soil | % | ≈250m |
| Soil Organic Carbon | 7 | Soil | g per kg | ≈250m |
| Soil Sand Content | 7 | Soil | mass fraction in % | ≈250m |
| Soil Saturated Water Content | 7 | Soil | % | ≈250m |
| Soil Silt Content | 7 | Soil | mass fraction in % | ≈250m |
| Soil pH | 7 | Soil | pH x 10 | ≈250m |
| Eastness | 3 | Topography | eastness index (-1 to 1) | 30 arcsec (≈900m at equator) |
| Elevation | 3 | Topography | meters | 30 arcsec (≈900m at equator) |
| Northness | 3 | Topography | northness index (-1 to 1) | 30 arcsec (≈900m at equator) |
| Roughness | 3 | Topography | (see reference) | 30 arcsec (≈900m at equator) |
| Slope | 3 | Topography | (see reference) | 30 arcsec (≈900m at equator) |

1. CHELSA [19,20]
2. CGIAR [21]
3. EarthEnv [22]
4. ESA CCI [23]
5. Obu *et al.* 2019
6. Fan *et al.* 2013
7. Soilgrids [26]

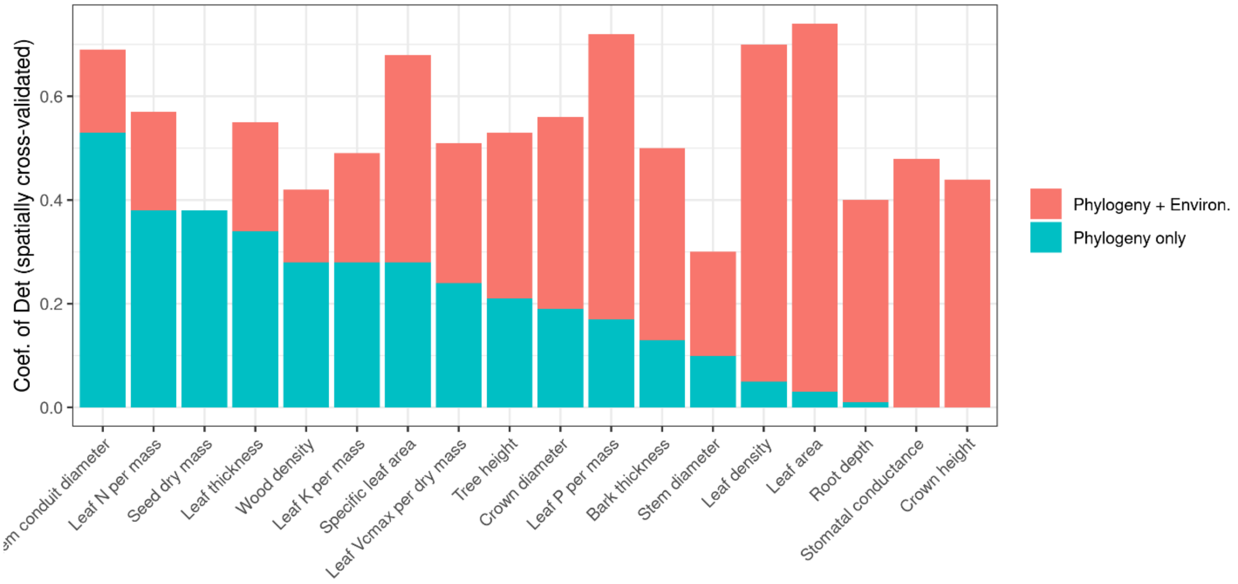


**Figure S1.** The performance of the models ($R^2_{VEcv}$) with and without environmental variables included alongside phylogenetic information.

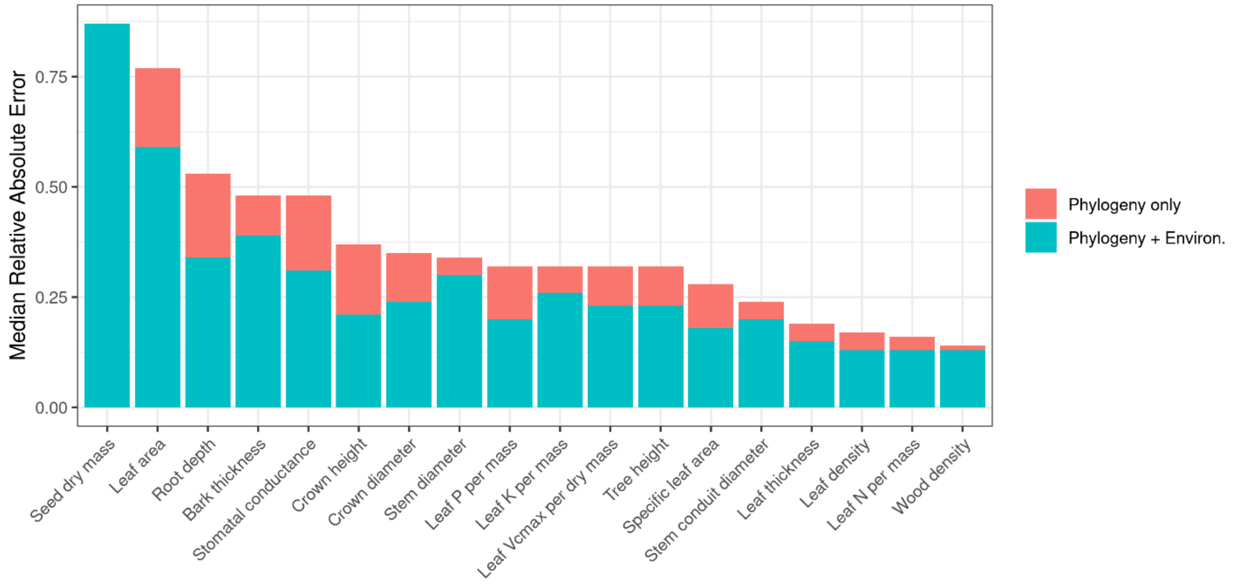


**Figure S2.** The performance of the models ($R^2_{VEcv}$) with and without phylogenetic and environmental variables included.

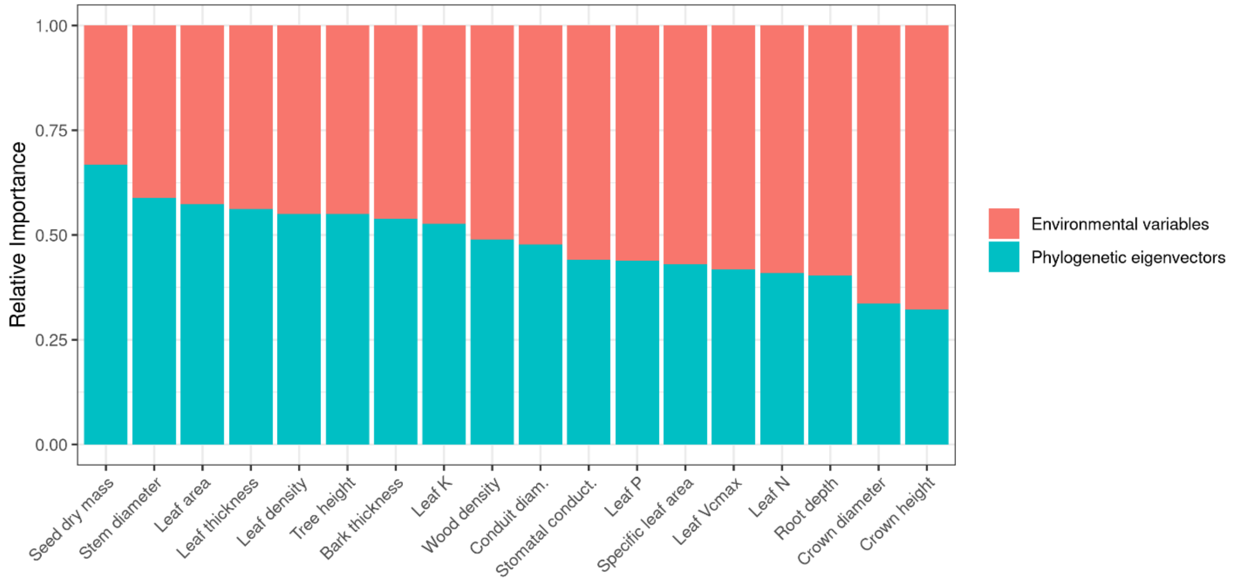


**Figure S3.** The relative importance (scaled to 1) attributable to environmental variables vs. phylogenetic eigenvectors as predictors of trait expression. R

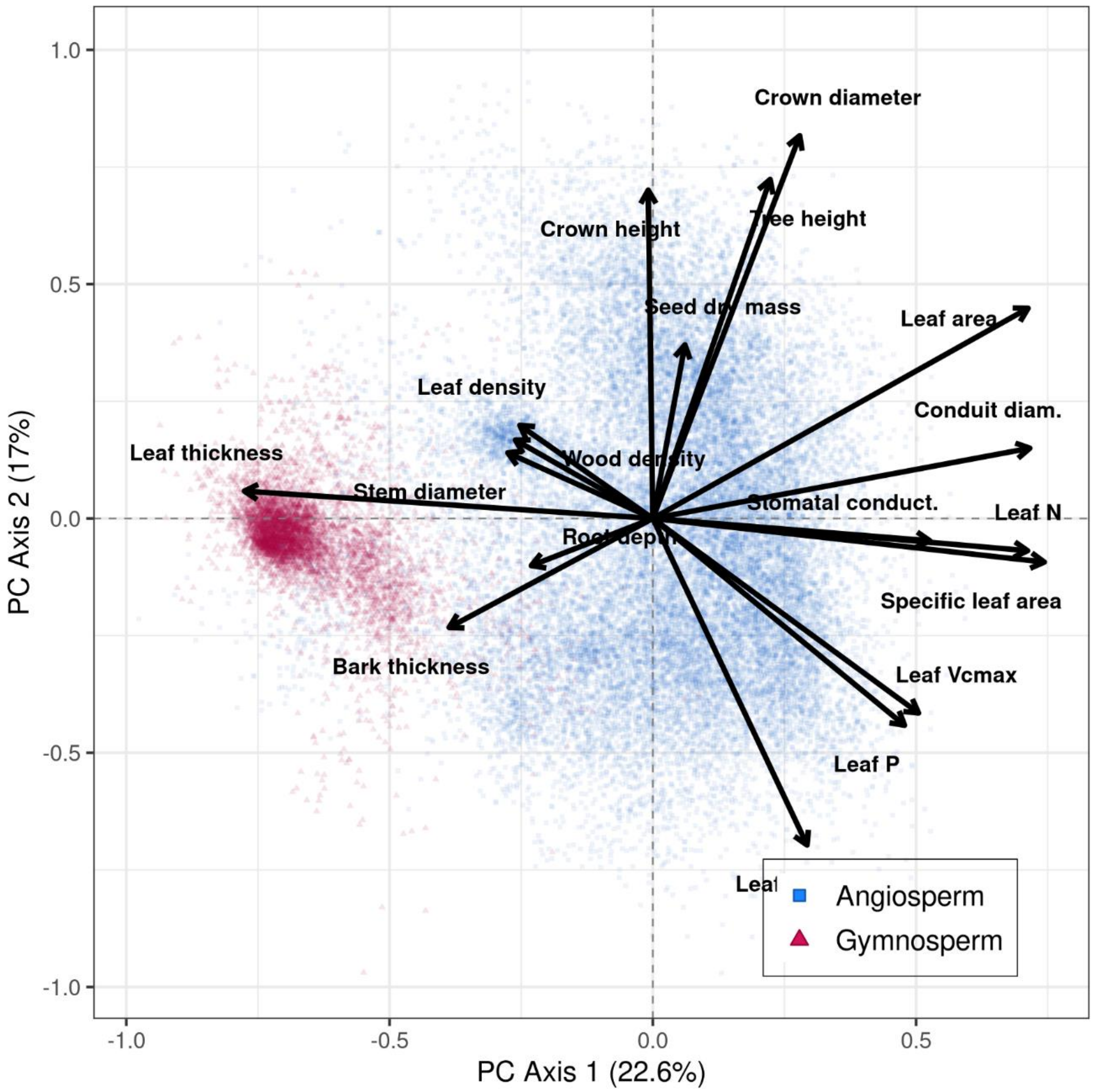


**Figure S4.** The full PCA for angiosperms + gymnosperms.

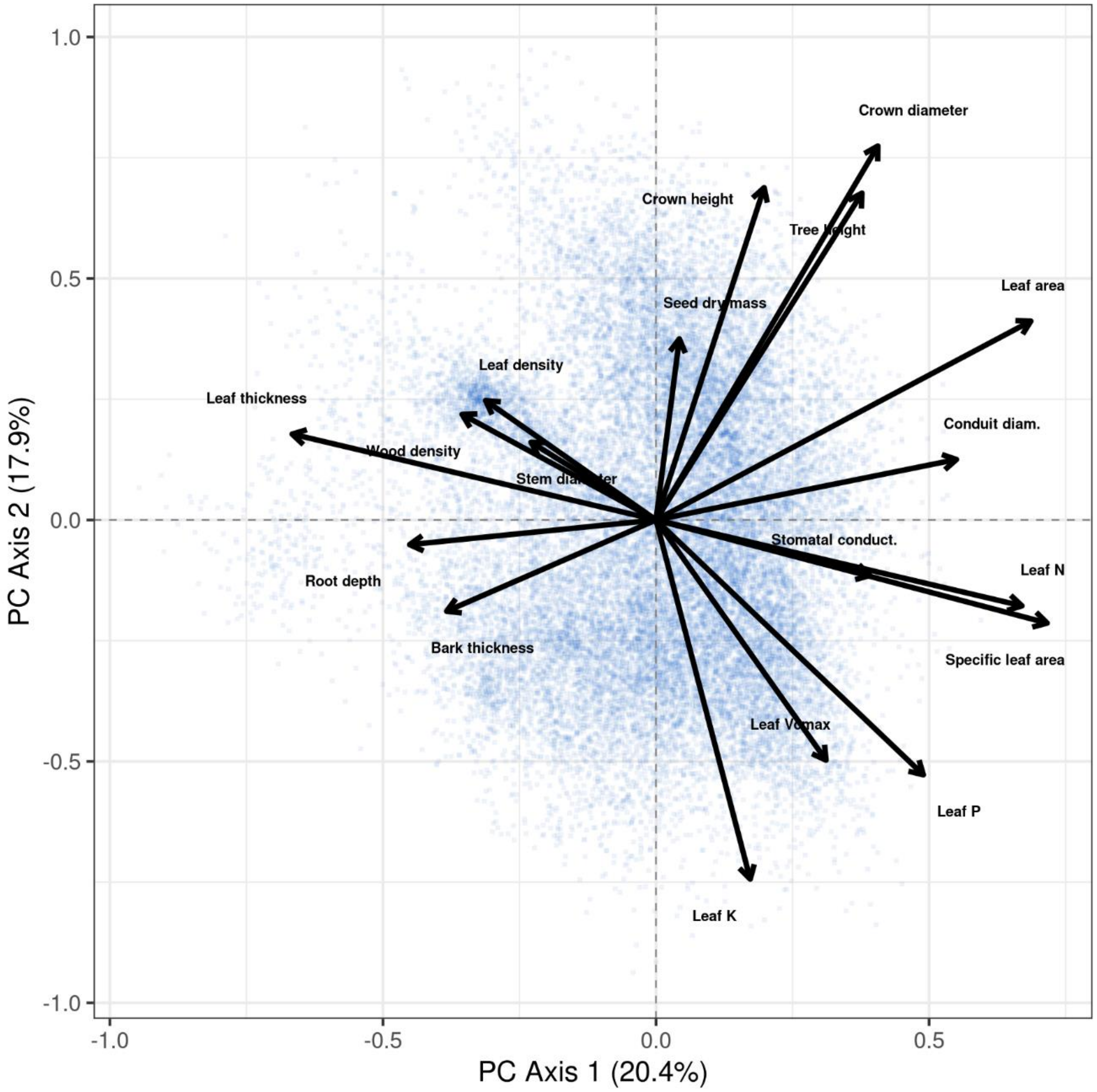


**Figure S5.** The full PCA for angiosperms only.

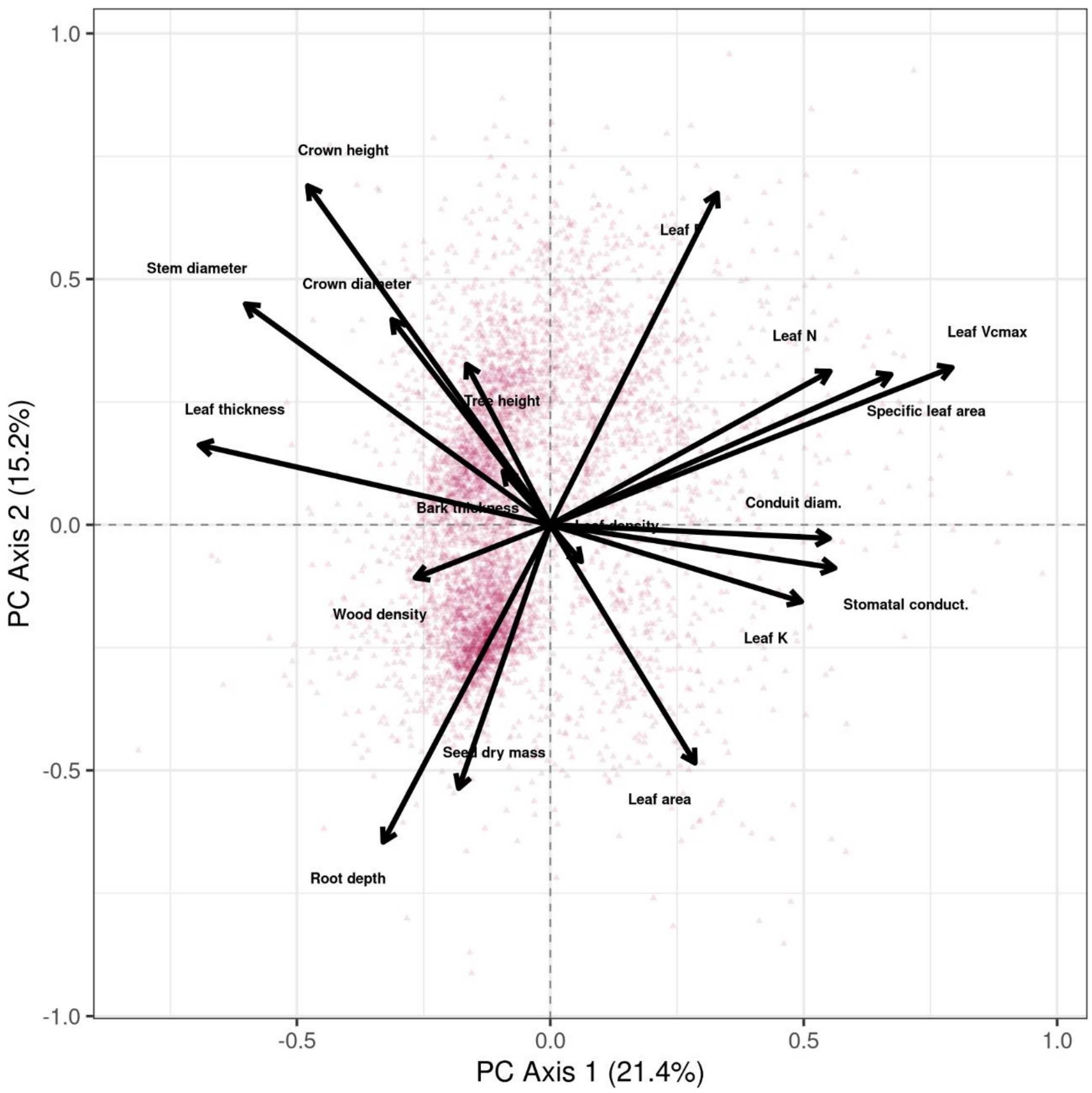


**Figure S6.** The full PCA for angiosperms only.

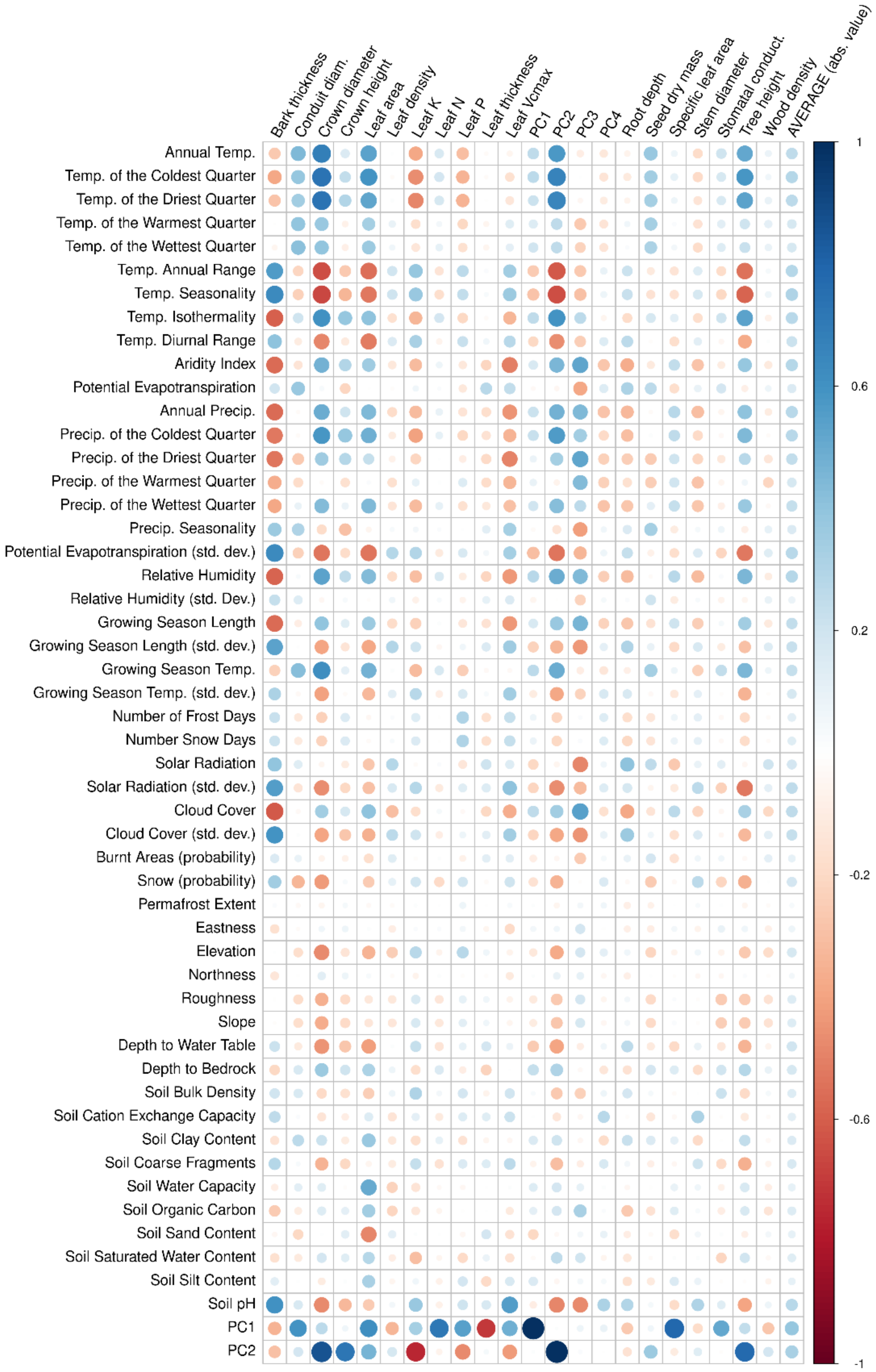


**Figure S7.** Spearman rank correlations between the 18 traits, the 50 environmental variables, and two primary trait PCA trade-offs.

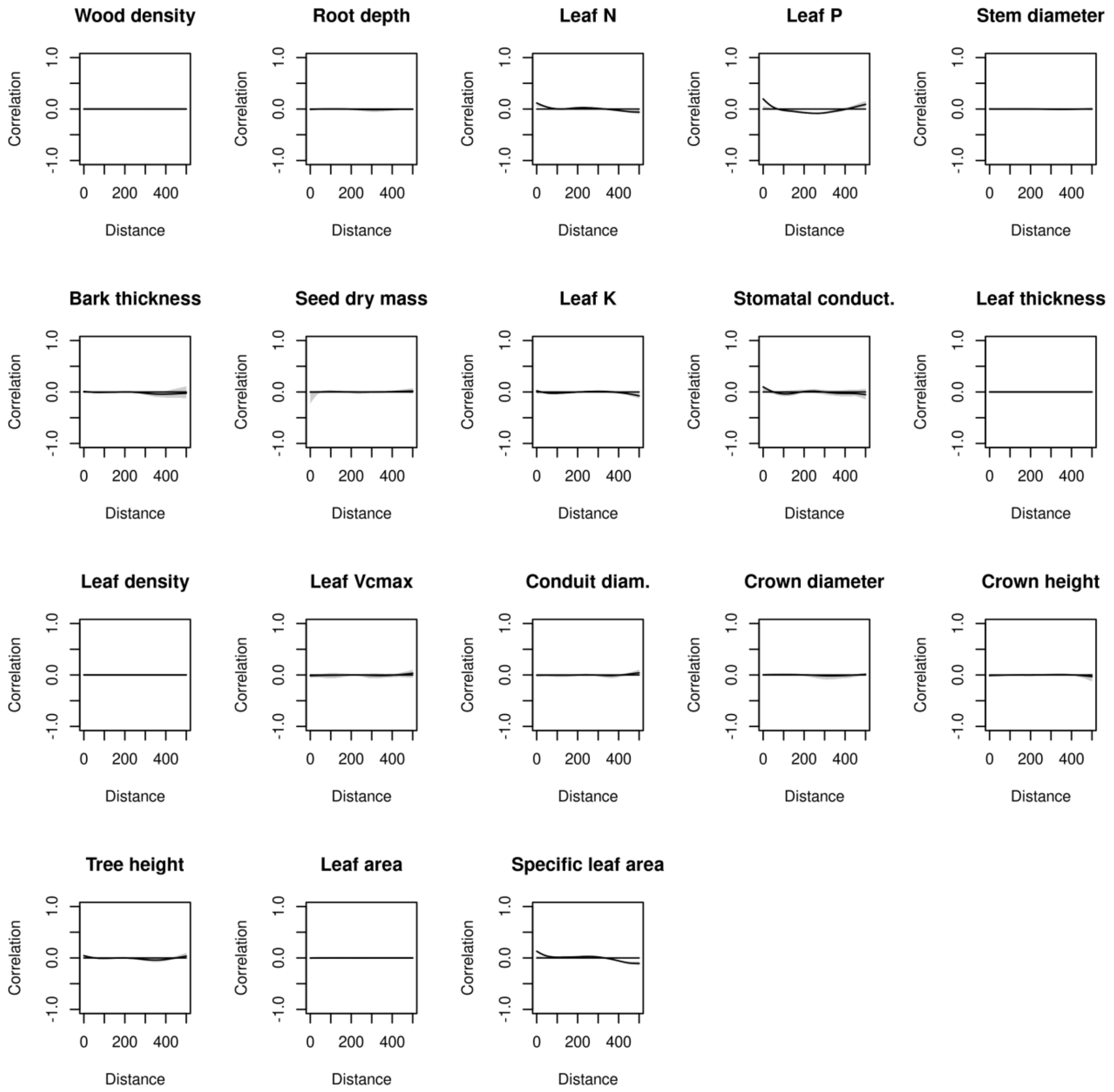


**Figure S8.** Residual spatial autocorrelation (Moran's I) for all 18 traits, assessed using standard linear regression models.

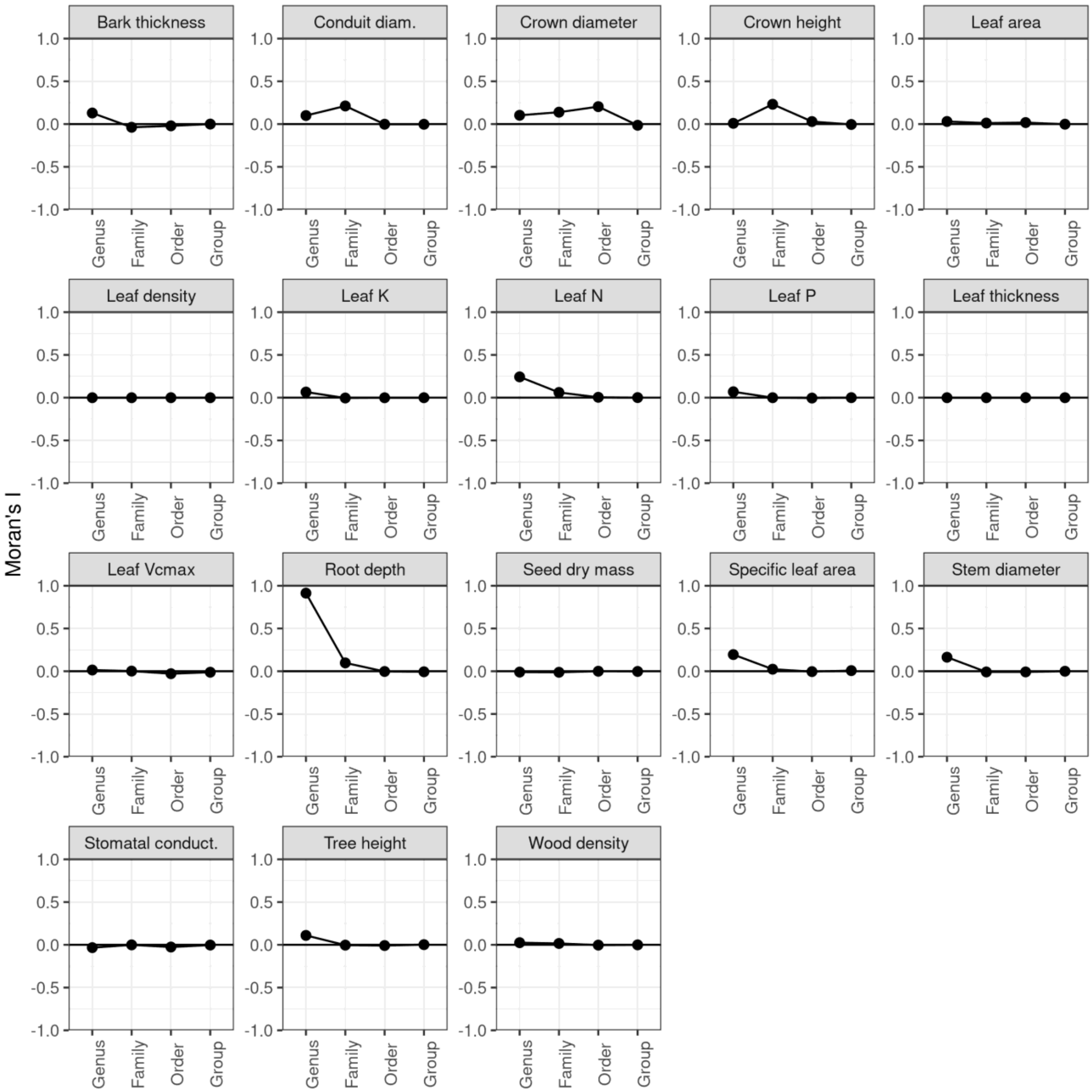


**Figure S9.** Residual taxonomic autocorrelation (Moran's I) for all 18 traits, assessed using standard linear regression models.

**Supplemental Data References**


1. Adriaenssens S. (2012). Dry deposition and canopy exchange for temperate tree species under high nitrogen deposition. PhD thesis, Ghent University, Ghent, Belgium, 209p.
2. Atkin OK (2015) Global variability in leaf respiration among plant functional types in relation to climate and leaf traits. New Phytologist DOI: 10.1111/nph.13253
3. Aubin, I., Messier, C., Gachet, S., Lawrence, K., McKenney, D., Arseneault, A., Bell, W., De Grandpré, L., Shipley, B., Ricard, J.P. and Munson, A.D., 2012. TOPIC-traits of plants in Canada. Natural Resources Canada, Canadian Forest Service, Sault Ste. Marie, Ontario. Online [URL] TOPIC website :http://cfs.cloud.nrcan.gc.ca/ctn/topic.php
4. Auger, S., Shipley, B. (2012). Interspecific and intraspecific trait variation along short environmental gradients in an old-growth temperate forest. Journal of Vegetation Science. DOI: 1111/j.1654-1103.2012.01473.x
5. Bahar, NHA, Ishida, FY, Weerasinghe, LK, Guerrieri, R, OSullivan, OS, Bloomfield, KJ, Asner, GP, Martin, RE, Lloyd, J, Malhi, Y, Phillips, OL, Meir, P, Salinas, N, Cosio, EG, Domingues, TF, Quesada, CA, Sinca, F, Escudero Vega, A, Zuloaga Ccorimanya, PP, del Aguila-Pasquel, J, Quispe Huaypar, K, Cuba Torres, I, Butrón Loayza, R, Pelaez Tapia, Y, Huaman Ovalle, J, Long, BM, Evans, JR, Atkin, OK (2017) Leaf-level photosynthetic capacity in lowland Amazonian and high-elevation Andean tropical moist forests of Peru. New Phytologist 214, 1002-1018.
6. Baraloto, C., C. E. T. Paine, L. Poorter, J. Beauchene, D. Bonal, A.-M. Domenach, B. Herault, S. Patino, J.-C. Roggy, and J. Chave. 2010. Decoupled leaf and stem economics in rainforest trees. Ecology Letters 13:1338-1347
7. Baruch, Z. & Goldstein, G. 1999. Leaf construction cost, nutrient concentration, and net CO2 assimilation of native and invasive species in Hawaii. Oecologia 121: 183-192
8. Blonder, B., Baldwin, B., Enquist, B.J., Robichaux, R.H. (2016) Variation and macroevolution in leaf functional traits in the Hawaiian silversword alliance (Asteraceae). Journal of Ecology 104:219-228 DOI: 10.1111/1365-2745.12497

25. Cornelissen, J. H. C., H. M. Quested, D. Gwynn-Jones, R. S. P. Van Logtestijn, M. A. H. De Beus, A. Kondratchuk, T. V. Callaghan, and R. Aerts. 2004. Leaf digestibility and litter

decomposability are related in a wide range of subarctic plant species and types. Functional Ecology 18:779-786.

26. Cornwell, W. K., R. Bhaskar, L. Sack, S. Cordell, and C. K. Lunch. 2007. Adjustment of structure and function of Hawaiian Metrosideros polymorpha at high vs. low precipitation. Functional Ecology 21:1063-1071.

27. Craine, J. M., A. J. Elmore, M. P. M. Aidar, M. Bustamante, T. E. Dawson, E. A. Hobbie, A. Kahmen, M. C. Mack, K. K. McLauchlan, A. Michelsen, G. B. Nardoto, L. H. Pardo, J. Penuelas, P. B. Reich, E. A. G. Schuur, W. D. Stock, P. H. Templer, R. A. Virginia, J. M. Welker, and I. J. Wright. 2009. Global patterns of foliar nitrogen isotopes and their relationships with climate, mycorrhizal fungi, foliar nutrient concentrations, and nitrogen availability. New Phytologist 183:980-992.

28. Craven, D., D. Braden, M. S. Ashton, G. P. Berlyn, M. Wishnie, and D. Dent. 2007. Between and within-site comparisons of structural and physiological characteristics and foliar nutrient content of 14 tree species at a wet, fertile site and a dry, infertile site in Panama. Forest Ecology and Management 238:335-346.

29. Díaz, S., J. G. Hodgson, K. Thompson, M. Cabido, J. H. C. Cornelissen, A. Jalili, G. Montserrat-Martí, J. P. Grime, F. Zarrinkamar, Y. Asri, S. R. Band, S. Basconcelo, P. Castro-Díez, G. Funes, B. Hamzehee, M. Khoshnevi, N. Pérez-Harguindeguy, M. C. Pérez-Rontomé, F. A. Shirvany, F. Vendramini, S. Yazdani, R. Abbas-Azimi, A. Bogaard, S. Boustani, M. Charles, M. Dehghan, L. de Torres-Espuny, V. Falczuk, J. Guerrero-Campo, A. Hynd, G. Jones, E. Kowsary, F. Kazemi-Saeed, M. Maestro-Martínez, A. Romo-Díez, S. Shaw, B. Siavash, P. Villar-Salvador, and M. R. Zak. 2004. The plant traits that drive ecosystems: Evidence from three continents. Journal of Vegetation Science 15:295-304.

30. Dahlin KM, Asner GP & CB Field (2013) Environmental and community controls on plant canopy chemistry in a Mediterranean-type ecosystem. Proceedings of the National Academy of Sciences USA. 110(17): 6895-6900

31. Dawson, S. K., Warton, D. I., Kingsford, R. T., Berney, P. , Keith, D. A., Catford, J. A. and Mori, A. (2017), Plant traits of propagule banks and standing vegetation reveal

flooding alleviates impacts of agriculture on wetland restoration. J Appl Ecol, 54: 1907-1918. doi:10.1111/1365-2664.12922

32. de Araujo, A.C., J. P. H. B. Ometto, A. J. Dolman, B. Kruijt, M. J. Waterloo and J. R. Ehleringer. 2011. LBA-ECO CD-02 C and N Isotopes in Leaves and Atmospheric CO2, Amazonas, Brazil. Data set. Available on-line [http://daac.ornl.gov ] from Oak Ridge National Laboratory Distributed Active Archive Center, Oak Ridge, Tennessee, U.S.A.

33. de Araujo, A.C., J.P.H.B. Ometto, A.J. Dolman, B. Kruijt, M.J. Waterloo and J.R. Ehleringer. 2012. LBA-ECO CD-02 C and N Isotopes in Leaves and Atmospheric CO2, Amazonas, Brazil. Data set. Available on-line [http://daac.ornl.gov ] from Oak Ridge National Laboratory Distributed Active Archive Center, Oak Ridge, Tennessee, U.S.A. http://dx.doi.org/10.3334/ORNLDAAC/1097

34. Domingues TF, Meir P, Feldpausch TR, et al. (2010) Co-limitation of photosynthetic capacity by nitrogen and phosphorus in West Africa woodlands. Plant, Cell & Environment (33): 959-980.

35. Falster DS, Remko A. Duursma, Masae I. Ishihara, Diego R. Barneche, Richard G. FitzJohn, Angelica Våhammar, Masahiro Aiba, Makoto Ando, Niels Anten, Michael J. Aspinwall, Jennifer L. Baltzer, Christopher Baraloto, Michael Battaglia, John J. Battles, Ben Bond-Lamberty, Michiel van Breugel, James Camac, Yves Claveau, Lluís Coll, Masako Dannoura, Sylvain Delagrange, Jean-Christophe Domec, Farrah Fatemi, Wang Feng, Veronica Gargaglione, Yoshiaki Goto, Akio Hagihara, Jefferson S. Hall, Steve Hamilton, Degi Harja, Tsutom Hiura, Robert Holdaway, Lindsay B. Hutley, Tomoaki Ichie, Eric J. Jokela, Anu Kantola, Jeff W. G. Kelly, Tanaka Kenzo, David King, Brian D. Kloeppel, Takashi Kohyama, Akira Komiyama, Jean-Paul Laclau, Christopher H. Lusk, Douglas A. Maguire, Guerric le Maire, Annikki Mäkelä, Lars Markesteijn, John Marshall, Katherine McCulloh, Itsuo Miyata, Karel Mokany, Shigeta Mori, Randall W. Myster, Masahiro Nagano, Shawna L. Naidu, Yann Nouvellon, Anthony P. OGrady, Kevin L. OHara, Toshiyuki Ohtsuka, Noriyuki Osada, Olusegun O. Osunkoya, Pablo Luis Peri, Any Mary Petritan, Lourens Poorter, Angelika Portsmuth, Catherine Potvin, Johannes Ransijn, Douglas Reid, Sabina C. Ribeiro, Scott D. Roberts, Rolando Rodríguez, Angela Saldaña-

Acosta, Ignacio Santa-Regina, Kaichiro Sasa, N. Galia Selaya, Stephen C. Sillett, Frank Sterck, Kentaro Takagi, Takeshi Tange, Hiroyuki Tanouchi, David Tissue, Toru Umehara, Hajime Utsugi, Matthew A. Vadeboncoeur, Fernando Valladares, Petteri Vanninen, Jian R. Wang, Elizabeth Wenk, Richard Williams, Fabiano de Aquino Ximenes, Atsushi Yamaba, Toshihiro Yamada, Takuo Yamakura, Ruth D. Yanai, and Robert A. York. 2015. BAAD: a biomass and allometry database for woody plants. Ecology 96:1445.http://dx.doi.org/10.1890/14-1889.1

41. Garnier, E., S. Lavorel, P. Ansquer, H. Castro, P. Cruz, J. Dolezal, O. Eriksson, C. Fortunel, H. Freitas, C. Golodets, K. Grigulis, C. Jouany, E. Kazakou, J. Kigel, M. Kleyer, V. Lehsten, J. Leps, T. Meier, R. Pakeman, M. Papadimitriou, V. P. Papanastasis, H. Quested, F. Quetier, M. Robson, C. Roumet, G. Rusch, C. Skarpe, M. Sternberg, J.-P. Theau, A. Thebault, D. Vile, and M. P. Zarovali. 2007. Assessing the effects of land-use change on plant traits, communities and ecosystem functioning in grasslands: A standardized

methodology and lessons from an application to 11 European sites. Annals of Botany 99:967-985.

42. Givnish T.J., R.A. Montgomery and G. Goldstein. 2004. Adaptive radiation of photosynthetic physiology in the Hawaiian lobeliads: light regimes, static light responses, and whole-plant compensation points. American Journal of Botany 91: 228-246

43. Gonzalez-Akre, E., McShea, W., Bourg, N., Anderson-Teixeira, K. 2015. Leaf traits data (SLA) for 56 woody species at the Smithsonian Conservation Biology Institute-ForestGEO Forest Dynamic Plot. Front Royal, Virginia. USA. [Data set]. Version 1.0.(www.try-db.org)

49. HIGUCHI, P.; SILVA, A.C. Araucaria Forest Database. 2013

50. Hoof, J., L. Sack, D. T. Webb, and E. T. Nilsen. 2008. Contrasting structure and function of pubescent and glabrous varieties of Hawaiian Metrosideros polymorpha (Myrtaceae) at high elevation. Biotropica 40:113-118.

51. Iversen CM, McCormack ML, Powell AS, Blackwood CB, Freschet GT, Kattge J, Roumet C, Stover DB, Soudzilovskaia NA, Valverde-Barrantes OJ, van Bodegom PM, Violle C

(2017) A global Fine-Root Ecology Database to address belowground challenges in plant ecology. New Phytologist. doi:10.1111/nph.14486.

52. Jennifer S. Powers and Peter Tiffin 2012 Plant functional type classifications in tropical dry forests in Costa Rica: leaf habit versus taxonomic approaches. Functional Ecology 2010, 24, 927-936 doi: 10.1111/j.1365-2435.2010.01701.x

53. Joseph, G.S., Seymour, C.L., Cumming, G.S., Cumming, D.H.M., & Mahlangu, Z. 2014. Termite mounds increase functional diversity of woody plants in African savannas. Ecosystems 17: 808-819.

54. Kattge, J., W. Knorr, T. Raddatz, and C. Wirth. 2009. Quantifying photosynthetic capacity and its relationship to leaf nitrogen content for global-scale terrestrial biosphere models. Global Change Biology 15:976-991.

55. Kichenin et al. 2013. Contrasting effects of plant inter- and intraspecific variation on community-level trait measures along an environmental gradient. Functional Ecology, in press.

56. Kleyer, M., R. M. Bekker, I. C. Knevel, J. P. Bakker, K. Thompson, M. Sonnenschein, P. Poschlod, J. M. van Groenendael, L. Klimes, J. Klimesova, S. Klotz, G. M. Rusch, Hermy, M. , D. Adriaens, G. Boedeltje, B. Bossuyt, A. Dannemann, P. Endels, L. Götzenberger, J. G. Hodgson, A.-K. Jackel, I. Kühn, D. Kunzmann, W. A. Ozinga, C. Römermann, M. Stadler, J. Schlegelmilch, H. J. Steendam, O. Tackenberg, B. Wilmann, J. H. C. Cornelissen, O. Eriksson, E. Garnier, and B. Peco. 2008. The LEDA Traitbase: a database of life-history traits of the Northwest European flora. Journal of Ecology 96:1266-1274.

62. Lhotsky B., Anikó Csecserits, Bence Kovács, Zoltán Botta-Dukát: New plant trait records of the Hungarian flora

63. Li, R., Zhu, S., Chen, H. Y. H., John, R., Zhou, G., Zhang, D., Zhang, Q. and Ye, Q. (2015), Are functional traits a good predictor of global change impacts on tree species abundance dynamics in a subtropical forest?. Ecol Lett, 18: 1181-1189. doi:10.1111/ele.12497

64. Lin Y-S, Medlyn BE, Duursma RA, Prentice IC, Wang H, Baig S, Eamus D, De Dios VR, Mitchell P, Ellsworth DS, De Beeck MO, Wallin G, Uddling J, Tarvainen L, Linderson M-L, Cernusak LA, Nippert JB, Ocheltree TW, Tissue DT, Martin-StPaul NK, Rogers A, Warren JM, De Angelis P, Hikosaka K, Han Q, Onoda Y, Gimeno TE, Barton CVM, Bennie J, Bonal D, Bosc A, Löw M, Macinins-Ng C, Rey A, Rowland L, Setterfield SA, Tausz-Posch S, Zaragoza-Castells J, Broadmeadow MSJ, Drake JE, Freeman M, Ghannoum O, Hutley LB, Kelly JW, Kikuzawa K, Kolari P, Koyama K, Limousin J-M, Meir P, Da Costa ACL, Mikkelsen TN, Salinas N, Sun W, Wingate L, (2015) Optimal stomatal behaviour around the world. Nature Climate Change 5(5): 459-464 DOI: 10.1038/NCLIMATE2550

65. Lukeš, P., Stenberg, P., Rautiainen, M., Mõttus, M., Vanhatalo, K.M. Optical properties of leaves and needles for boreal tree species in Europe (2013) Remote Sensing Letters, 4 (7), pp. 667-676

66. Lusk, C. H., Kaneko, T., Grierson, E. and Clearwater, M. (2013) Correlates of tree species sorting along a temperature gradient in New Zealand rain forests: seedling functional traits, growth and shade tolerance. Journal of Ecology, 101: 1531-1541.

67. Maire V, Ian J. Wright, I. Colin Prentice, Niels H. Batjes, Radika Bhaskar, Peter M. van Bodegom, Will K. Cornwell, David Ellsworth, Ülo Niinemets, Alejandro Ordoñez, Peter

B. Reich, Louis S. Santiago (2015). Global soil and climate effects on leaf photosynthetic traits and rates. Global Ecology and Biogeography 24(6): 706-717. Maire V, Wright IJ, Prentice IC, Batjes NH, Bhaskar R, van Bodegom PM, Cornwell WK, Ellsworth D, Niinemets Ü, Ordoñez A, Reich PB, Santiago LS (2015) Data from: Global effects of soil and climate on leaf photosynthetic traits and rates. Dryad Digital Repository. http://dx.doi.org/10.5061/dryad.j42m7

96. Rodrigues, A.V.; Bones, F.L.V.; Schneiders, A.; Oliveira, L.Z.; Vibrans, A.C.; Gasper, A.L. Plant Trait Dataset for Tree-Like Growth Forms Species of the Subtropical Atlantic Rain Forest in Brazil. Data 2018, 3, 16.

110. Souza K, Higuchi P, Silva Ac, Schimalski Mb, Loebens R, Buzzi Junior F, Souza Cc, Rodrigues Junior Lc, Walter Ff, Missio Ff, Dalla Rosa A. (2017) Partição de nicho por

grupos funcionais de espécies arbóreas em uma floresta subtropical. Rodriguésia [online] vol.68, n.4, pp.1165-1175. ISSN 0370-6583. http://dx.doi.org/10.1590/2175-7860201768401

111. Spasojevic, M. J., Turner, B. L., and Myers, J. A. (2016) When does intraspecific trait variation contribute to functional beta?diversity? J Ecol, 104: 487-496. doi:10.1111/1365-2745.12518

112. Swaine, E. K. 2007. Ecological and evolutionary drivers of plant community assembly in a Bornean rain forest. PhD Thesis, University of Aberdeen, Aberdeen.

113. Takkis, K. 2014. Changes in plant species richness and population performance in response to habitat loss and fragmentation. Dissertationes Biologicae Universitatis Tartuensis 255, 2014-04-07. Available from: http://hdl.handle.net/10062/39546

114. Tamir Klein, Giovanni Di Matteo, Eyal Rotenberg, Shabtai Cohen, Dan Yakir (2012) Differential ecophysiological response of a major Mediterranean pine species across a climatic gradient. Tree Physiology 33 (1): 26-36. doi: 10.1093/treephys/tps116

115. Thuiller W - Traits of European Alpine Flora - Wilfried Thuiller - OriginAlps Project - Centre National de la Recherche Scientifique

126. Wirth, C. and J. W. Lichstein. 2009. The Imprint of Species Turnover on Old-Growth Forest Carbon Balances - Insights From a Trait-Based Model of Forest Dynamics.

Pages 81-113 in C. Wirth, G. Gleixner, and M. Heimann, editors. Old-Growth Forests: Function, Fate and Value. Springer, New York, Berlin, Heidelberg.
